# The WT1-BASP1 complex is required to maintain the differentiated state of taste receptor cells

**DOI:** 10.1101/460816

**Authors:** Yankun Gao, Debarghya Dutta Banik, Stefan G. E. Roberts, Kathryn F. Medler

## Abstract

The WT1-BASP1 complex is a transcriptional repressor that is involved in cell differentiation; however, the mechanisms underlying its effects are not well-characterized *in vivo*. In the peripheral taste system which endogenously expresses these proteins, BASP1 co-expression with WT1 begins at the end of development, suggesting a role for the WT1-BASP1 complex in terminal cell differentiation within this system. Using a conditional BASP1 mouse, we demonstrate that BASP1 is critical to maintain the differentiated state of adult cells by inhibiting the activation of WT1-dependent target genes. Our results uncover a central role for the WT1-BASP1 complex in maintaining cell differentiation.

## Introduction

The WT1 transcription factor plays a critical role in the development and maintenance of multiple organs and tissues (Hastie 2017). In particular, WT1 null mice display complete agenesis of the kidneys, gonads, adrenal glands and spleen. WT1 is also needed for tissue maintenance in the adult, with these sites having some overlap with developmental targets as well as additional organs (Chau et al. 2011). WT1 can either drive cell proliferation or promote differentiation, but the mechanisms involved in this dichotomy are not clear (Toska and Roberts 2014; Hastie 2017).

WT1 often acts in concert with a transcriptional cofactor, BASP1. BASP1 binding switches the function of WT1 from an activator to a repressor (Toska and Roberts 2014) and regulates the ability of WT1 to control differentiation in several model cell lines including kidney podocyte cells (Green et al. 2009), epicardial cells (Essafi et al. 2011) and blood cells (Goodfellow et al. 2011). Recent work has also shown that in the absence of BASP1, WT1 has an important role in maintaining multipotency. BASP1 blocks this function and is associated with driving iPSCs to differentiate (Blanchard et al. 2017). Thus BASP1 is a critical regulator of WT1 function.

WT1 null mice also have developmental defects in several sensory tissues including the retinal ganglion cells (Wagner et al. 2002), olfactory epithelia (Wagner et al. 2005) and, as shown by us, peripheral taste cells (Gao et al. 2014). A unique feature of peripheral taste cells is that they are continuously replaced throughout an organism’s lifetime (Barlow and Klein 2015) which generates a need for constant remodelling of these cells. We also found that WT1 is expressed in adult taste cells but its role is currently unknown.

Taste receptor cells originate from Keratin 14 (Krt14+) expressing progenitor cells that become either non-taste epithelium or post-mitotic precursors that express sonic hedgehog (Shh+). These post-mitotic Shh+ cells further differentiate into functional taste cells that express Keratin 8 (Krt8). Krt8 is highly expressed in the mature taste cells which are located in taste buds present in the oral cavity. These cells are divided into one of three groups (type I, II or III) which are based on their physiological function as well as the expression of specific markers and anatomical features (Liu et al. 2013; Barlow and Klein 2015).

The current understanding of the taste cell renewal process is far from complete. It is clear that both the Shh and Wnt/β-catenin signaling pathways regulate the specification of taste cell fate and are required for taste cell differentiation (Castillo et al. 2014; Gaillard et al. 2015; Gaillard et al. 2017). However, the underlying mechanisms regulating Wnt and Shh signaling in adult taste cells during this process are still unknown. The goal of this study was to analyze the role of BASP1 within taste cell renewal. We find that deletion of BASP1 in differentiated cells leads to their reduced function, a loss of several cell type markers normally found in mature cells, and the upregulation of WT1 target genes that are primarily expressed in the progenitor cells. Our findings reveal that the WT1-BASP1 complex plays a central role in the maintenance of the differentiated state in this system.

## Results and Discussion

Our previous work identified a key role for WT1 in the development of the peripheral taste system, specifically the circumvallate (CV) papillae (Gao et al. 2014). The CV papillae is an epithelial specialization located on the back of the tongue which houses hundreds of taste buds. Since BASP1 is a critical transcriptional cofactor for WT1, we sought to determine if it acts with WT1 during the development of this papillae type. We analyzed BASP1 expression at different developmental stages during CV formation using double label immunohistochemistry with either Krt8+ (a marker of the developing taste cells; Figure 1A), or GAP43 (a marker that identifies the gustatory nerve which will innervate the emerging taste cells; Figure 1B). At E12.5 and E15.5, BASP1 is present in the developing gustatory nerve (GAP43+) but is not co-expressed with Krt8 in the placode that is developing into the CV papillae. By P7, BASP1 is expressed in both the Krt8+ cells and GAP43+ gustatory nerve. This expression pattern is maintained in the adult taste system. Thus, unlike WT1 (Gao et al. 2014), BASP1 is not expressed during CV development but is switched on around birth in these cells. BASP1 expression is maintained in the adult taste buds while its expression in the gustatory nerve is reduced after birth. Adult mouse taste cells express WT1 (Gao et al. 2014) and we find that it shows a large degree of colocalization with BASP1 (Figure 2A). To identify the specific cell types that express WT1 and BASP1,βperformed immunohistochemistry using unique markers for each of the identified cell types. NTPDase2 is a marker specific to the type I cells and has strong co-localization with WT1 expression (Figure 2B) but does not co-localize with BASP1 (Figure 2C). In the type II (PLCβ2+) and III (NCAM+) cells both WT1 and BASP1 are expressed. Thus, WT1 is expressed in all three types of differentiated taste cells while BASP1 expression is restricted to the type II and III cells. Type II and III cells are the transducers of taste stimuli while type I cells are thought to function as support cells similar to glial cells in the central nervous system (Finger 2000).

**Figure 1:**
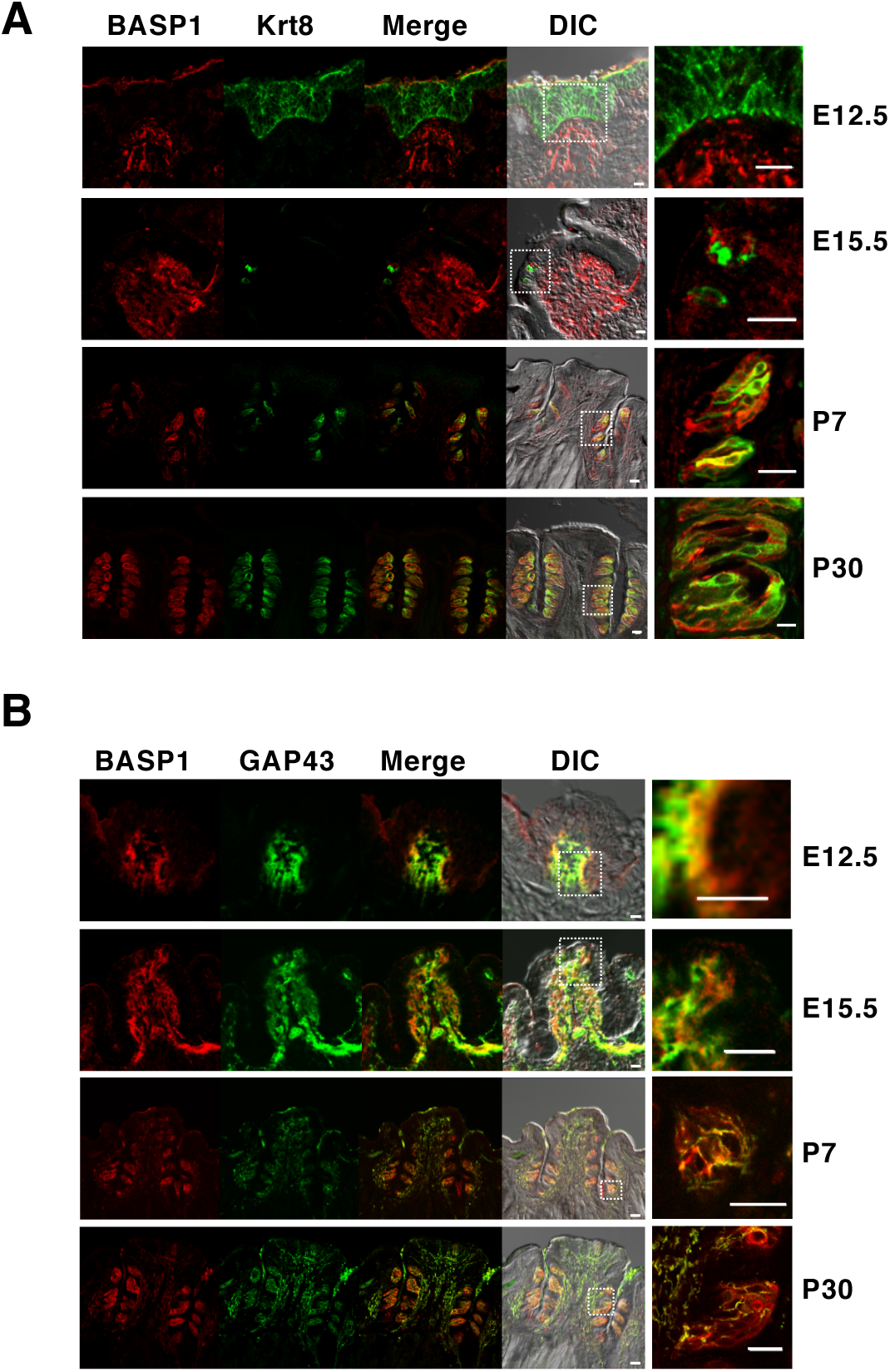
BASP1 is expressed at the terminal stages of CV development. **(A)** Immunohistochemistry of CV at different stages of development (indicated at right). BASP1 is red and Krt8 is green and merged image is shown. Higher magnification of the merged image (boxed section) is shown at right. **(B)** As in part A except that BASP1 (red) and GAP43 (green) were detected. Scale bars=10μM.

**Figure 2:**
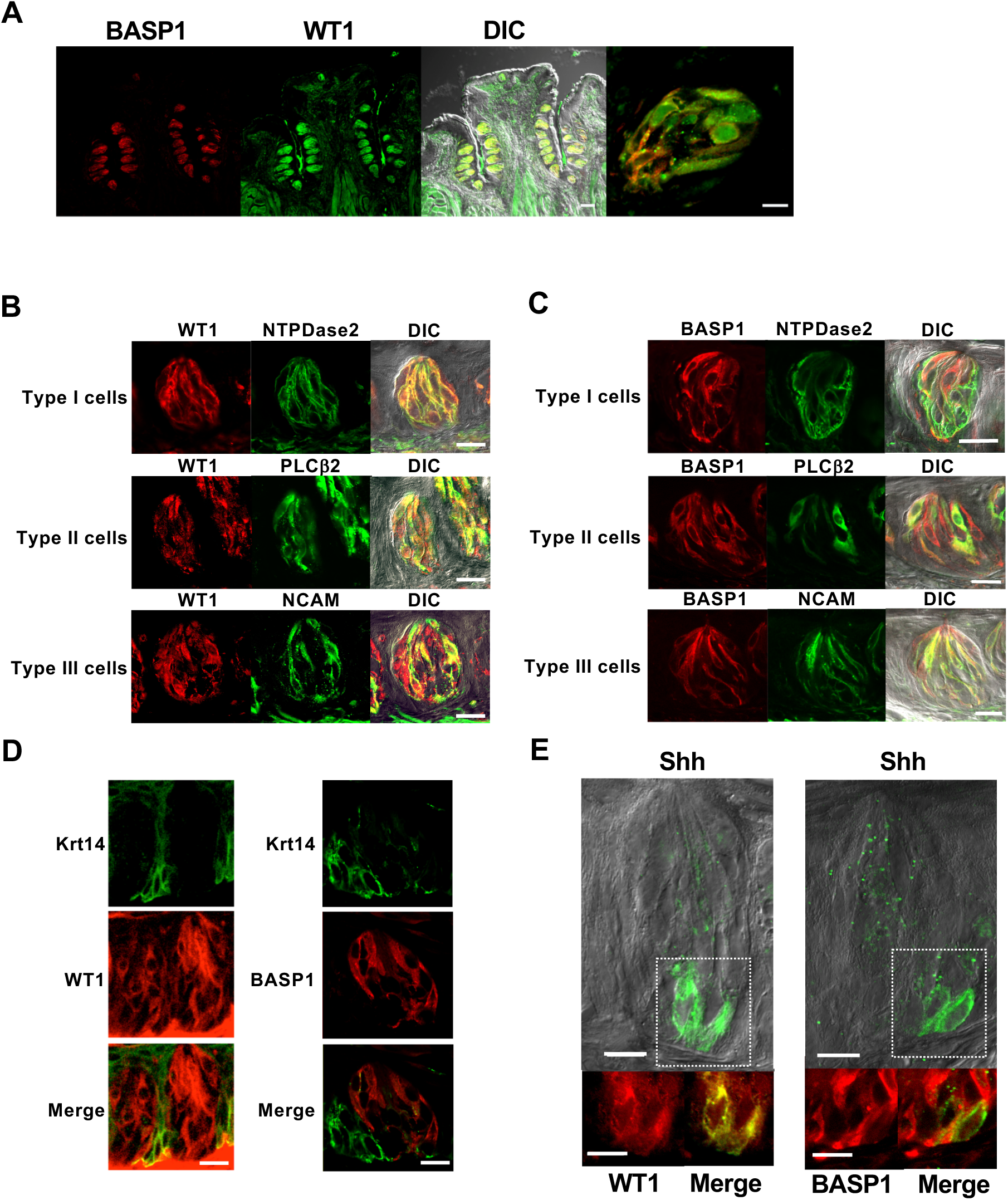
WT1 and BASP1 are expressed in overlapping cell types of the adult CV. **(A)**WT1 and BASP1 are expressed in adult taste cells Immunohistochemistry of CV papillae to detect WT1 (green) and BASP1 (red) and merged image. Higher magnification of the merged image showing an individual taste bud (boxed section) is shown at right. **(B)** Immunohistochemistry of CV taste buds to detect WT1 (red) and either NTPDase2 (green; type I cells), PLCβ2 (green; type II cells) or NCAM (green; type III cells). Merged images are shown at right. **(C)** As in part B except that BASP1 (red) was detected instead of WT1. **(D)** Left panels; Immunohistochemistry of CV taste buds to detect WT1 (red) and Krt14 (green). Merged image is shown at bottom. Right panels; Immunohistochemistry of CV taste buds to detect BASP1 (red) and Shh (green). Merged image is shown at bottom right. **(E)** As in part D except that Shh (green) was detected instead of Krt14. Scale bars=10 μ M.

We also found that WT1, but not BASP1, is expressed in the progenitor (Krt14+) cells (Figure 2D). This finding suggests that WT1 is functioning very early in the cell renewal process during the proliferation and formation of new taste cells. This is consistent with WT1’s function in the CV during embryonic development (Gao et al. 2014) as well as its role in iPSCs (Blanchard et al. 2017). WT1 is also found in the post-mitotic progenitor (Shh+) cells that will differentiate into the mature cells, while nuclear BASP1 is expressed in a subset of the Shh+ cells (n=7 of 27 cells). This raises the possibility that BASP1 expression is initiated as the Shh+ cells undertake the final transition to terminal differentiation (Figure 2E). Taken together, our results demonstrate that WT1 is expressed in both the progenitors and differentiated taste cells while BASP1 expression is generally limited to the differentiated cells.

Our next experiments measured the effects of BASP1 on taste cell function. We crossed a floxed BASP1 mouse (Supplementary Figure 1) with a Krt8-Cre-ER mouse to delete BASP1 in the differentiated type I, II and III taste cells. Mice were treated with tamoxifen for 8 days, sacrificed a week later, and immunohistochemistry was performed to confirm knockout of BASP1 in the taste buds (Figure 3A). RNA was also prepared from isolated taste cells to confirm that BASP1 mRNA in taste cells was reduced (Figure 3A, at right). The specific deletion of the BASP1 gene in Krt8-expressing cells (Krt8-BASP1-KO; KO) did not lead to any obvious change in the taste bud morphology compared to controls (CTL) and the number of taste buds in the CV papillae of KO mice was not significantly different from CTL mice (Figure 3B). Furthermore, the expression levels of Krt8 in the taste buds was not altered in the KO mice (Figure 3C).

**Figure 3:**
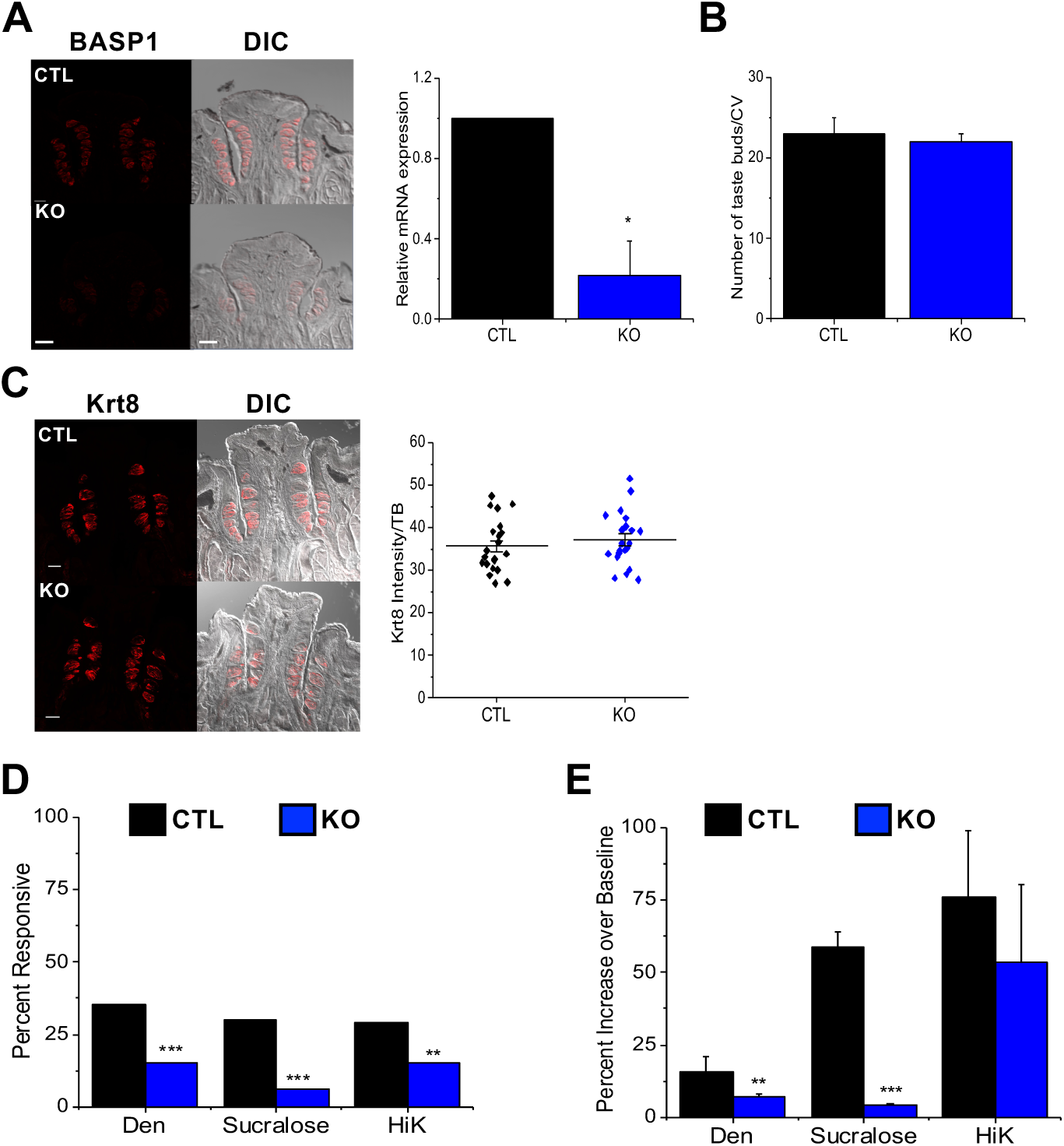
Knockout of BASP1 in the adult CV affects the stimulus responses of taste cells. **(A)** Immunohistochemistry of CV papillae to detect BASP1 (red) in control (CTL) and Krt8-BASP1-CRE (KO) mice treated with tamoxifen. Scale bar=50μM. At right qPCR to determine BASP1 expression relative to GAPDH in RNA isolated from CTL (black bar) and KO (blue bar, KO) mice (*,p<0.05). **(B)** Taste buds were stained with DAPI and the number of cells per bud were counted in CTL and KO mice. Mean with standard deviation are reported. **(C)** As in part A except that staining was with anti-Krt8. At right is quantitation of immunoreactivity per taste bud using Image J. Horizontal bars represent mean average intensity. **(D)** Chi-square analysis compared the overall number of responsive taste cells in CTL and KO mice. The percentage of responsive taste cells is shown (*** p < 0.001 and **, p<0.01) for 5mM denatonium, 20mM sucralose and 50mM potassium chloride (HiK). **(E)** The response amplitudes (percent increase over baseline) for the responsive cells from part D were analyzed and compared for the CTL and KO mice using a Student’s t test (***p < 0.001 and**, p<0.01). Representative traces are shown in Supplemental Figure 2.

To evaluate if the loss of BASP1 affected cellular function, we isolated taste cells and compared the taste-evoked calcium responses between KO and CTL mice using live cell imaging. Two taste stimuli were used to activate type II cells: sweet (20mM sucralose) and bitter (5mM denatonium benzoate, Den). Potassium chloride (50mM, HiK) was used to depolarize type III taste cells and activate voltage gated calcium channels (VGCCs). There was a significant reduction in the number of responsive cells for all three stimuli in KO mice compared to CTLs (Figure 3D). Moreover, the remaining calcium signals for denatonium and sucralose were significantly reduced in amplitude (percent increase over baseline) in KO mice compared to CTLs (Figure 3E). Representative traces are shown in Supplementary Figure 2. Taken together, these data reveal that the ability of the taste cells to respond to stimuli was significantly impaired when BASP1 expression was ablated.

Our results so far demonstrate that the expression of BASP1 in differentiated taste cells is required for their normal function and that the absence of BASP1 significantly reduced their ability to respond to stimuli. We then measured the effect of BASP1 deletion on the specific markers for each cell type. Deletion of BASP1 (Figure 4A) caused a significant increase in the expression of the type I cell marker, NTPDase2 (Figure 4B). We did not detect BASP1 in the type I cells of CTL mice (Figure 2C), suggesting either that a low level of BASP1 in type I cells normally acts to modulate NTPDase2 expression or that deletion of BASP1 in other Krt8+ cells affects the properties of type I cells. Conversely, loss of BASP1 in Krt8+ cells caused the reduction of both PLCβ2 (type II cell marker; Figure 4C) and NCAM (type III cell marker; Figure 4D) expression. The reduction in PLCβ2 and NCAM expression was not due to a decrease in the overall number of type II and III cells (Figure 4E), but was instead due to the lower expression of these proteins in the cells (Figure 4C and D, graphs). Not all type II and type III functional proteins were affected by BASP1 deletion as gustducin (type II marker) and pgp9.5 (type III marker) expression levels were maintained in the KO mice (Supplementary Figure 3). The reduced expression of PLCβ2 and NCAM suggests a loss of cell specialization and is consistent with the impaired responses of the taste cells that we observed (Figure 3). Taken together, these data (Figures 3 and 4) suggest that loss of BASP1 in the differentiated taste cells leads to a significant alteration in their properties.

**Figure 4:**
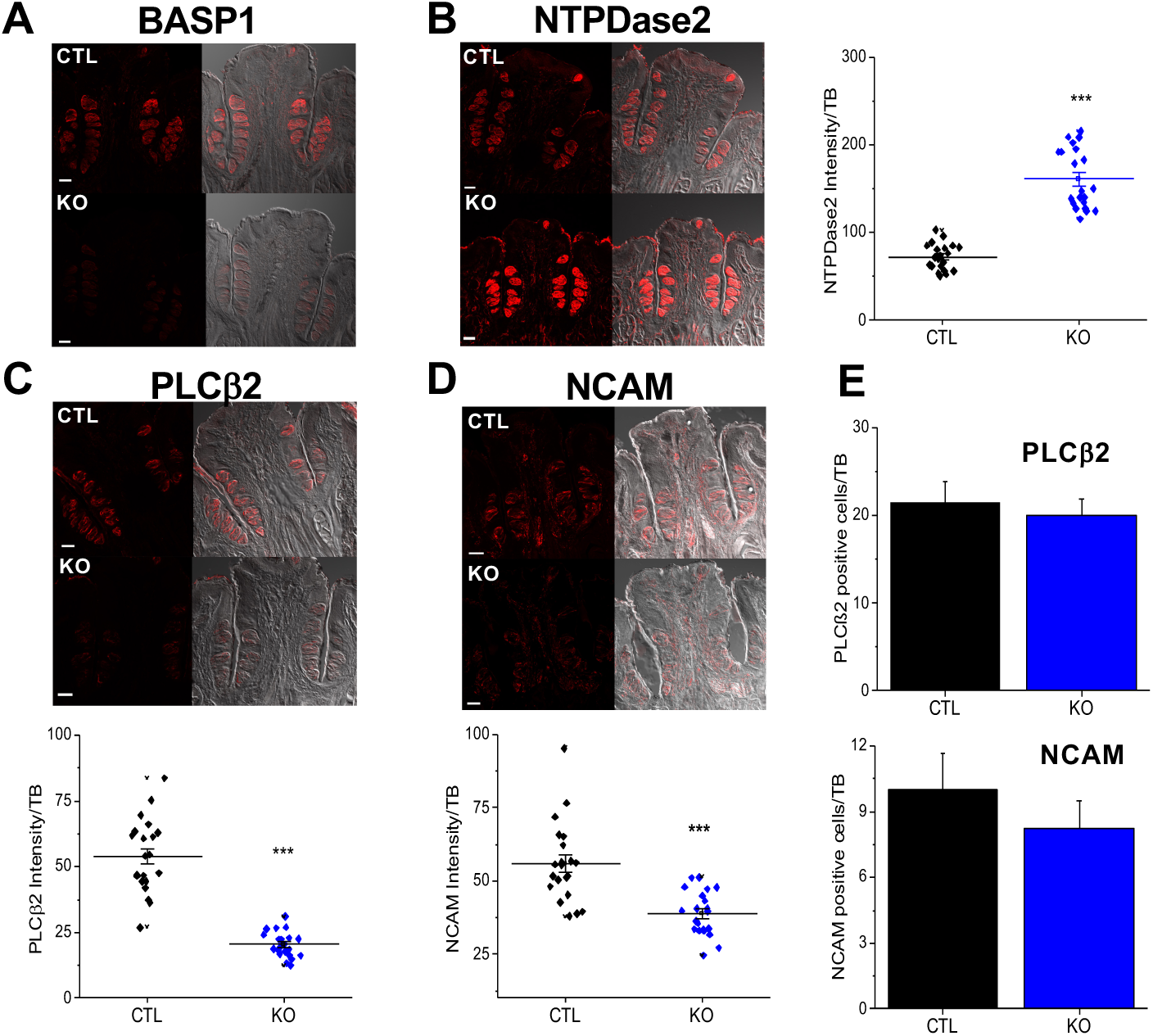
Knockout of BASP1 in the adult CV leads to a disruption of type II and type III cells markers. **(A)** Immunohistochemistry of CV taste buds using anti-BASP1 (red) in CTL and KO mice that had been treated with tamoxifen. Scale bar=50μM. **(B)** As in part A except that NTPDase2 (type I marker) was tested. A plot of signal intensity per taste bud is shown at right. Horizontal bars represent mean intensity (*** p<0.001). **(C)** As in part A except that anti-PLCβ2 (type II marker) was used. A plot of signal intensity per taste bud is shown below. Horizontal bars represent mean intensity (***, p<0.001). **(D)** As in part A except that NCAM labelling (type III marker) was measured. A plot of signal intensity per taste bud is shown below. Horizontal bars represent mean intensity (***p<0.001). **(E)** The number of PLCβ2-(above) and NCAM-(below) positive cells per taste bud in CTL and KO cells is shown. Error bars are standard deviation of the mean.

Our previous work identified the LEF1 and PTCH1 genes as direct targets for WT1 regulation during the development of the CV papillae (Gao et al. 2014). These genes encode components of the Wnt and Shh signalling pathways, respectively, that are important during the development of the peripheral taste system. These signalling pathways also play a central role during the early stages of the cell renewal process in the precursor cells (Krt14+, Shh+ cells) but are generally switched off in the fully differentiated taste cells (Barlow and Klein 2015). To determine if LEF1 and PTCH1 expression were affected by the loss of BASP1 in the differentiated taste cells, we prepared RNA from isolated taste cells for either KO or CTL mice. qPCR was used to measure the levels of LEF1 and PTCH1 mRNA relative to control GAPDH mRNA (Figure 5A) in the KO and CTL cells. Deletion of BASP1 in the Krt8-positive taste cells led to a significant increase in both LEF1 and PTCH1 expression compared to CTL.

**Figure 5:**
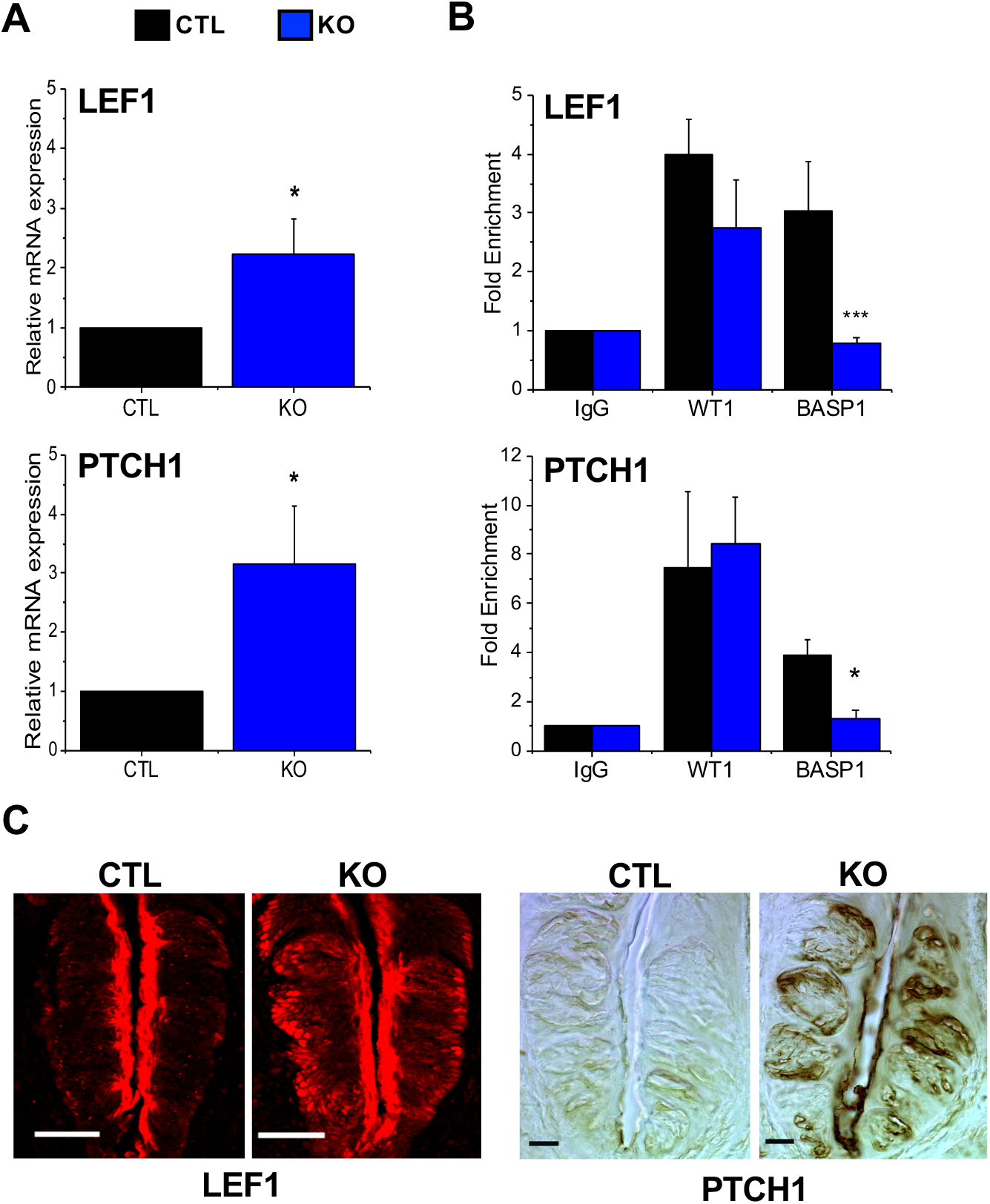
The WT1-BASP1 complex represses LEF1 and PTCH1 expression in the differentiated cells of the CV. **(A)** Quantitative PCR (qPCR) was used to detect LEF1 (above) and PTCH1 (below) expression relative to GAPDH in mRNA isolated from taste cells of CTL and KO mice (*, p<0.05). **(B)** ChIP analysis of isolated taste cells from adult mice. WT1, BASP1 or control (IgG) antibodies were used. After normalization to input DNA, fold enrichment at the LEF1 (top) or PTCH1 (bottom) promoter regions over a control genomic region is shown in CTL and KO mice (*, p<0.05, ***, p<0.001). **(C)** Immunohistochemical analysis of LEF1 expression in the CV of CTL or KO mice (left panels). Expression levels of PTCH1 in the CV of CTL or KO mice (right panels). Scale bars= 20μM.

We then determined if WT1 and BASP1 directly localize to the promoter regions of the LEF1 and PTCH1 genes in these cells. Taste cells were isolated from the CV papillae of either CTL or KO mice and subjected to Chromatin Immunoprecipitation (ChIP) using WT1, BASP1, or control (IgG) antibodies. The results are expressed as fold enrichment at the LEF1 and PTCH1 promoters over a control genomic region (Figure 5B). While WT1 was localized at the LEF1 and PTCH1 promoters in both CTL and KO cells, BASP1 was only bound to the promoters of the LEF1 and PTCH1 genes in CTL cells and its binding was lost in the KO cells. Based on these data, we conclude that the WT1-BASP1 complex is directly repressing the expression of LEF1 and PTCH1 in the differentiated taste cells. When BASP1 is lost, the expression of both LEF1 and PTCH1 is significantly upregulated.

Immunohistochemical analysis evaluated the levels of LEF1 and PTCH1 protein expression in the CV of KO mice compared to CTLs (Figure 5C). LEF1 and PTCH1 are normally expressed at a low level in the mature cells and have higher expression levels in the undifferentiated progenitor cells (Miura et al. 2001; Gaillard et al. 2017). Loss of BASP1 in the differentiated taste cells (Krt8+) caused an upregulation in the expression of both LEF1 and PTCH1 throughout the CV papillae when compared to CTLs (Figure 5C). Taken together, the results in Figure 5 demonstrate that WT1-BASP1 acts as a transcriptional repressor complex in taste cells and that it normally inhibits the expression of Wnt and Shh signaling components in the differentiated cells.

Our previous work demonstrated that WT1 regulates the expression of LEF1 and PTCH1 genes during the embryonic development of the peripheral taste system, from very early stages through birth (Gao et al. 2014). We have now shown that BASP1 is not expressed during development of these cells but is expressed in the differentiated cells shortly after birth. WT1, but not BASP1, is expressed in the progenitor K14+ cells which constitutively express LEF1 and PTCH1. In the K14+ cells, these signalling pathways direct the differentiation pathway to Shh+ cells which further differentiate into the functional taste cells (Kapsimali and Barlow 2013; Barlow and Klein 2015; Gaillard et al. 2015). The selective expression of BASP1 in a subset of Shh+ cells suggests that BASP1 binding is needed to drive the Shh+ cells to differentiate into either type II or type III cells while WT1 alone is sufficient to drive differentiation of type I cells. One possibility is that BASP1 binding to WT1 in Shh+ cells causes the upregulation of gene expression pathways that are important for the specific functions of type II and III cells.

Our results show that the sustained presence of the WT1-BASP1 complex is required to maintain the taste cells in their differentiated state. Such a mechanism is consistent with the recently reported roles of WT1 and BASP1 in stem cells in which WT1 promotes multipotency while the WT1-BASP1 complex maintains the differentiated state and inhibits the induction of pluripotency (Blanchard et al. 2017). Several *in vitro* studies have demonstrated that BASP1 converts WT1 from a transcriptional activator to a repressor and that this switch allows the WT1/BASP1 complex to regulate the differentiated state of the cell (Toska and Roberts 2014). Our data are the first demonstration that this dual-control mechanism of the WT1/BASP1 complex is critical for the differentiated state of a system *in vivo*.

## Methods

### Animals

Floxed BASP1 mice were generated in a C57BL/6 background by Cyagen BioSciences (Supplementary Figure 1). Animals were cared for in compliance with the University at Buffalo Animal Care and Use Committee. BASP1fl^+/-^ mice were mated with Krt8-Cre/ERT2^+^ mice (017947, The Jackson Laboratory) to obtain BASP1fl^+/-^Krt8-Cre/ERT2^+^ mice. These offspring were bred to obtain BASP1fl^+^/^+^Krt8-Cre/ERT2^+^ mice which were used as BASP1-KO group. The BASP1fl^-/-^;Krt8-Cre/ERT2^+^ or BASP1fl^+/+^;Krt8-Cre/ERT2^-^ littermates were used as wild type CTLs. PCR primers used to genotype the conditional BASP1 knockout allele: 5’-TGCCCTGCCTGCAGGTCAAT-3’ (Forward); 5’-CCGAGTCTGTACAAAAGCCACC-3’ (Reverse). Tamoxifen was dissolved in corn oil mixed with ethanol (9:1 by volume) to a stock concentration of 20mg/ml as previously described (Miura et al. 2014). BASP1-KO and CTL mice were gavaged with tamoxifen (100mg/Kg body weight) every 24 hours for 8 consecutive days. Mice were sacrificed and tissues were collected 7 days after the last gavage. Primers used to genotype the Cre induced BASP1-KO allele: 5’-CGAGAGATTTGTGATGTATGATGAGCAG-3’ (F); 5’-CCGAGTCTGTACAAAAGCCACC −3’ (R). Taste cells were collected and calcium imaging performed as described in (Hacker et al. 2008; Maliphol et al. 2013; Dutta Banik et al. 2018).

### Immunohistochemistry

Fluorescent immunohistochemistry was performed as described in (Gao et al. 2014) except for experiments with LEF1 antibodies in which tongue sections were fixed for 10 seconds in 4% paraformaldehyde/0.1 M phosphate buffer (PB, pH 7.2) at room temperature. Fluorescence intensity was quantitated using Image J as previously described (Dutta Banik et al. 2018). PTCH1 immunohistochemistry was as follows; Mice were anesthetized with sodium pentobarbital (40 mg/kg; Patterson Veterinary) and then transcardially perfused with a 0.025% heparin solution in 1% sodium nitrite, followed by a 4% paraformaldehyde solution in 0.1 M phosphate buffer (pH 7.2). Following perfusion, the tongues were removed and placed into 4% paraformaldehyde for 2 h, followed by 4 °C overnight cryoprotection in 20% sucrose. On the following day, tongues were frozen in O.C.T. Compound (Sakura Finertek USA) and 12μm sections of the CV papillae were cut. Slides were incubated in 3% hydrogen peroxide blocking reagent for 5 minutes. After washing, the sections were incubated in serum blocking reagent, avidin blocking reagent and biotin blocking reagent for 15 minutes each. Sections were then incubated in PTCH1 or control antibodies for 2 hr at RT before an overnight incubation at 4°C. On the next day, sections were washed three times for 15 min each in buffer and incubated with biotinylated secondary antibody for 60 minutes, followed by 3 × 15 min washes in buffer. Sections were then incubated in HSS-HRP for 30 minutes, followed by 3 × 2min washes in buffer. Binding of primary antibody to the sections was visualized using a diaminobenzidine tetrahydrochloride (DAB) chromagen (R&D Biosystems).

Anti-Shh (sc-365112) and anti-gustducin (sc-395) were from Santa Cruz, anti-Krt8 from Developmental Studies Hybridoma Bank, anti-Lef1 from Signal Transduction (C12A5), anti-GAP43 from Aves (GAP43), anti-NTPDase2 from Ectonucleotidases-ab.com (CD39L1), anti-PLCβ2 from GeneTex (GTX133765), anti-NCAM (AB5032), anti-PGP9.5 (AB108986) and anti-Ptch1 (AB53715) were from Abcam, and anti-Krt14 from Biolegend (905301). WT1 and BASP1 antibodies were raised against a peptide (WT1) or recombinant protein (BASP1) by Pacific Immunology then purified by affinity chromatography. Antibody specificity was verified as described in (Toska et al. 2014). All experiments were performed on a minimum of three mice.

### RNA and ChIP analysis

Total RNA was extracted from taste cells using the Nucleospin RNA XS kit (Clontech). Unamplified total RNA was DNase treated and then reverse transcribed using the Bio-Rad cDNA synthesis kit. Primers for analysis of GAPDH, LEF1 and PTCH1 are described in (Gao et al. 2014). Samples were run in triplicate and at least three biological repeats were performed for each experiment. ChIP assays were performed as described before (Wang et al. 2010) using primers for LEF1, PTCH1 and control regions described in (Gao et al. 2014). Datasets are an average of three biological repeats with standard deviation and significance calculated by Students t-test.

## Acknowledgements

This work was funded by the NIH National Institute of General Medical Sciences [1R01GM098609 to K.F.M. and S.G.E.R.]. We thank Alan Siegel and the UB North Campus Imaging Facility funded by NSF-MRI Grant DBI 0923133 for the confocal images.

## Author contributions

K.F.M. and S.G.E.R. conceived the study, designed the experiments, analyzed the data and prepared the manuscript. Y.G. performed most of the experiments and D.D.B. performed the PTCH1 immunohistochemistry. Y.G. and D.D.B. analyzed the data and edited the manuscript.

---

**Supplementary Figure. 1.**
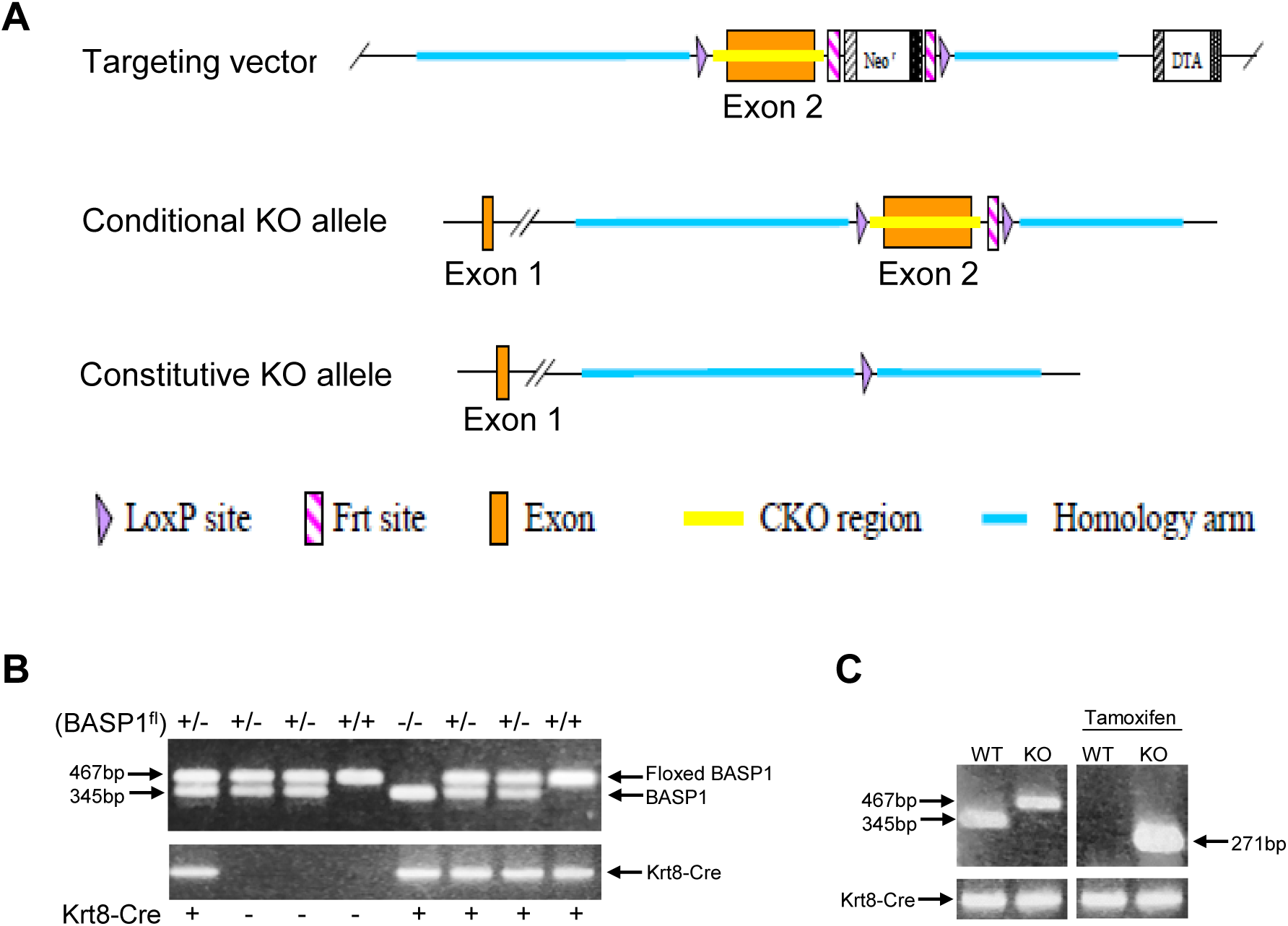
Strategy for producing a floxed BASP1 mouse. The mouse Basp1 gene (GenBank accession number: NM_027395.2, Ensembl: ENSMUSG00000045763) is located on mouse chromosome 15 and contains two exons. **(A)** Since all of the coding region for BASP1 is located solely in exon 2, it was targeted as the conditional knock out region. Deletion of exon 2 results in the total loss of Basp1 protein. To engineer the targeting vector, homology arms and CKO region were generated by PCR using BAC clone RP23-209B5 or RP23-334N16 from the C57BL/6J library as template. In the targeting vector, the Neo cassette was flanked by Frt sites, and CKO region was flanked by LoxP sites. DTA was used for negative selection. The conditional KO allele was obtained after Flp-mediated recombination and the constitutive KO allele is then obtained after Cre-mediated recombination. C57BL/6 ES cells were used for gene targeting. **(B)** Representative genotyping results from 8 mice with BASP1 and Krt8-CRE genotypes indicated. The upper gel is a compilation of homozygous floxed mice, heterozygous floxed mice and wild type (non-floxed) mice. The floxed band is present at 467bp while the wild type allele is 367bp. The lower gel identifies mice containing Krt8-Cre. **(C)** Prior to tamoxifen treatment, wild type and KO mice have either the 467 bp (floxed mouse) or the 345 bp band (wild type). After tamoxifen treatment, primers specific to the resulting floxed site that that is deleted of BASP1 produces a 271 bp band.

**Supplementary Figure 2:**
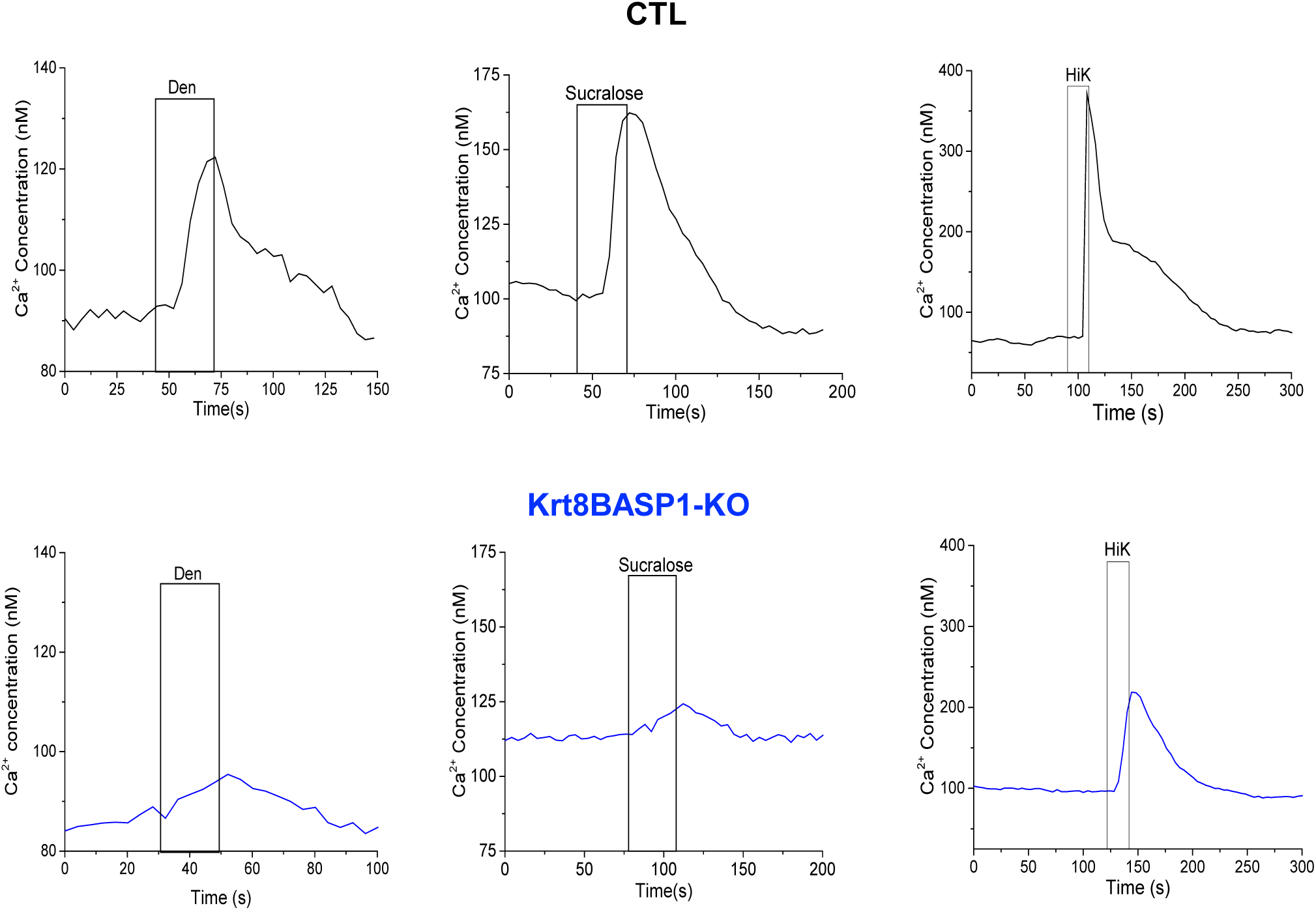
Reduced taste evoked response of CV cells in the absence of BASP1. Isolated taste cells from control mice (CTL) or Krt8-BASP1-KO mice were exposed to different taste stimuli and the resulting calcium response was measured using Fura2-AM in live cell imaging. Representative plots for each stimulus are shown.

**Supplementary Figure 3:**
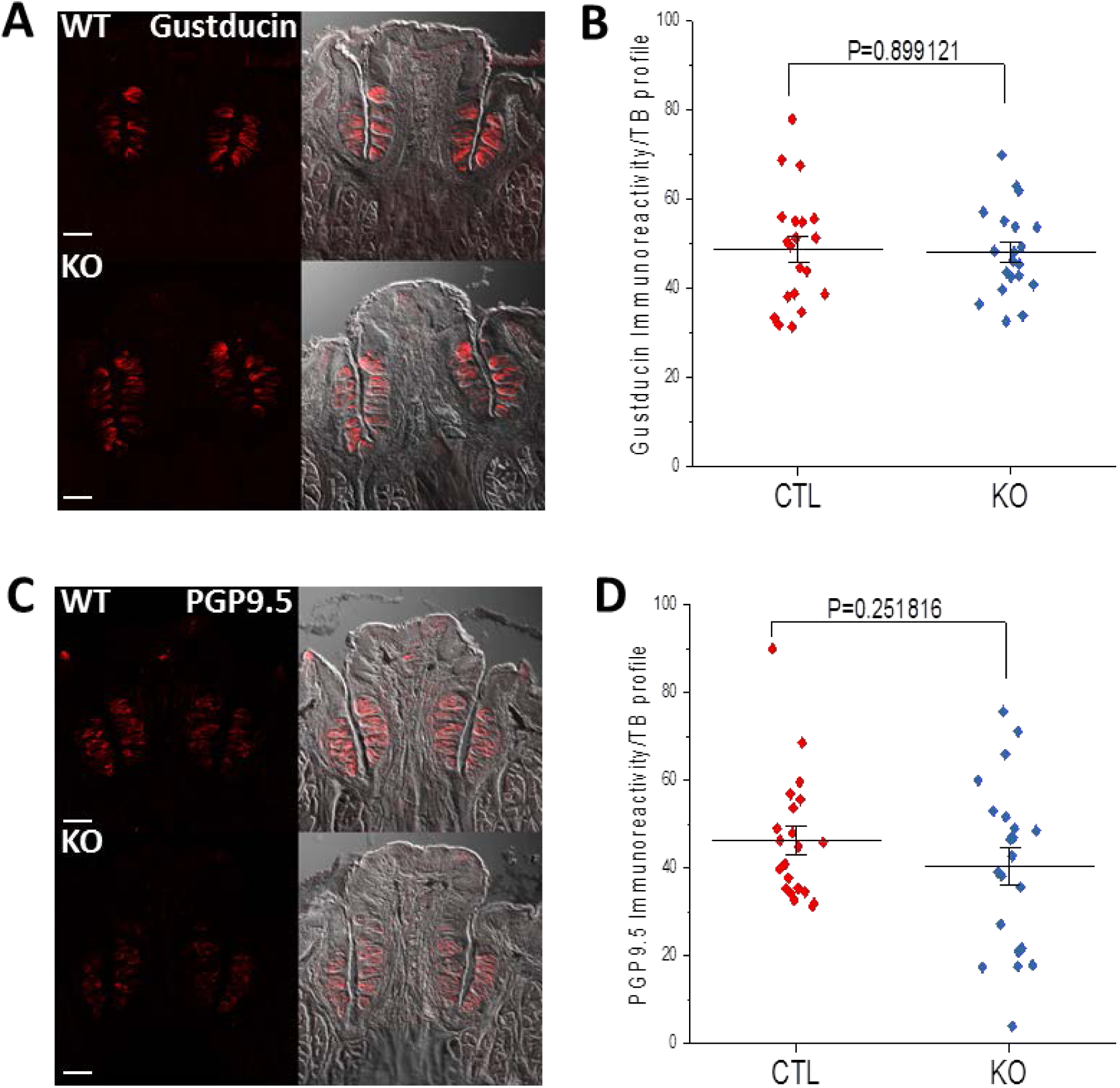
(A) Immunohistochemical analysis of CV taste buds to detect gustducin (red) in control (WT) mice and tamoxifen treated Krt8-BASP1-CRE (KO) mice. Scale bar=10μM. (B) A plot of gustducin signal intensity per taste bud in CTL and KO mice is shown. Horizontal bars represent mean intensity. (C) Immunohistochemistry of CV taste buds to detect PGP9.5 (red) in WT and KO mice treated with tamoxifen. Scale bar=10μM. (B) A plot of PGP9.5 signal intensity per taste bud in (CTL and KO mice) is shown. Horizontal bars represent mean intensity.

